# Local and global changes in cell density induce reorganisation of 3D packing in a proliferating epithelium

**DOI:** 10.1101/2024.02.08.579268

**Authors:** Vanessa Barone, Antonio Tagua, Jesus Á. Andrés-San Román, Amro Hamdoun, Juan Garrido-García, Deirdre C. Lyons, Luis M. Escudero

## Abstract

Tissue morphogenesis is intimately linked to the changes in shape and organisation of individual cells. In curved epithelia, cells can intercalate along their own apicobasal axes adopting a shape named “scutoid” that allows energy minimization in the tissue. Although several geometric and biophysical factors have been associated with this 3D reorganisation, the dynamic changes underlying scutoid formation in 3D epithelial packing remain poorly understood. Here we use live-imaging of the sea star embryo coupled with deep learning-based segmentation, to dissect the relative contributions of cell density, tissue compaction, and cell proliferation on epithelial architecture. We find that tissue compaction, which naturally occurs in the embryo, is necessary for the appearance of scutoids. Physical compression experiments identify cell density as the factor promoting scutoid formation at a global level. Finally, the comparison of the developing embryo with computational models indicates that the increase in the proportion of scutoids is directly associated with cell divisions. Our results suggest that apico-basal intercalations appearing just after mitosis may help accommodate the new cells within the tissue. We propose that proliferation in a compact epithelium induces 3D cell rearrangements during development.

**Summary statement:** The study uses sea star embryogenesis as a model of a proliferating epithelium to highlight how cell division induces 3D cell rearrangements during development.

## INTRODUCTION

Animal embryonic development is often driven by morphogenesis of epithelial tissues that form lumens, giving rise to hollow structures such as tubes or cysts that are the basis for further organogenesis (Guillot and Lecuit, 2013; Lecuit and Lenne, 2007, Navis and Bagnat, 2015). During these processes, epithelia become curved while maintaining their barrier function (Davidson, 2012; Pearl et al., 2017), which means that cell shape is adjusted so that the epithelium remains sealed and no openings or fractures occur (Gibson et al., 2006; Sánchez-Gutiérrez et al., 2016). Therefore, transformations in epithelial packing, i.e. the way in which epithelial cells are arranged in three dimensions, are crucial to the coupling of morphogenesis and function in curved epithelia (Gómez-Gálvez et al., 2021a; Lemke and Nelson, 2021).

In the example of monolayered epithelia, it has been traditionally assumed that cells have a prism shape and that curvature is achieved by prisms turning into frusta (i.e. truncated pyramids) (Davidson, 2012; Lecuit and Lenne, 2007; Pearl et al., 2017). Cells that are prisms or frusta have the same set of neighbours on their apical and basal sides (Schneider and Eberly, 2002). However, this is not always the case, as apico-basal intercalations can occur, i.e. neighbour exchanges that occur not in time, but in space, along the apicobasal axis of a cell - also known as apico-basal topological transition 1 (AB-T1) (Gómez-Gálvez et al. 2021; Gómez-Gálvez et al. 2018; Rupprecht et al. 2017; Sanchez-Corrales et al. 2018). Whenever an AB-T1 occurs, the four cells involved in the transition are no longer shaped as prisms or frusta; instead, they adopt another configuration, called scutoid, i.e. having different sets of neighbouring cells on their apical and basal surfaces (Gómez-Gálvez et al., 2022, 2021a, 2018; Lemke and Nelson, 2021; Lou et al., 2023; Mughal et al., 2018). AB-T1s occur frequently in curved epithelia (Honda et al., 2008; Sun et al., 2017; Xu et al., 2016), especially in tubular monolayered epithelia (Gómez-Gálvez et al. 2022; Gómez-Gálvez et al. 2018), allowing energetically favourable packing of cells in 3D (Gómez-Gálvez et al. 2022; Gómez-Gálvez et al. 2018). Based on this principle of energy minimization, AB-T1s are expected to form in any epithelium where the stresses acting on the apical and basal surfaces are anisotropic (Lou et al., 2023). This is the case in tissues with high curvature anisotropies, e.g. ovoids and tubes (Gómez-Gálvez et al., 2018; Mughal et al., 2018), or in areas with a large gradient of curvature where cells are laterally tilted (Lou et al., 2023; Rupprecht et al., 2017). In contrast, previous studies also suggest that geometrical cues should not induce AB-T1s in flat epithelia or in spherical epithelia, where the curvature is isotropic (do Carmo, 1976; Gómez-Gálvez et al., 2018; Lou et al., 2023).

Despite those predictions, AB-T1s *have* in fact been observed in curved epithelia without pronounced anisotropy and in higher frequencies than expected considering only tissue geometry. Therefore, it has been proposed that scutoid formation may also be due to cell and tissue dynamics producing differential strain on the apical and basal surfaces of epithelial tissues (Gómez-Gálvez et al., 2018; Lou et al., 2023; Rupprecht et al., 2017). For example, cell divisions may temporarily alter the balance of forces acting on epithelial cells (Gómez et al., 2021; Ragkousi et al., 2017; Ragkousi and Gibson, 2014). Similarly, the strains acting on tissues undergoing morphogenesis may be different from those predicted according to geometry alone, for instance due to active cell movements within the tissue (Gómez-Gálvez et al., 2018; Sanchez-Corrales et al., 2018; Sun et al., 2017) or to pressure from outer structures (Lou et al., 2023; Rupprecht et al., 2017). Determining whether dynamic processes underlie scutoid formation, requires following those processes in time. Therefore, high-resolution time-lapse imaging coupled with precise image segmentation is pivotal (Arganda-Carreras et al., 2017; Falk et al., 2019; Haberl et al., 2018; Lee et al., 2020; Wolny et al., 2020). This approach allows one to obtain realistic information about cell conformations in 3D, how they change over time and how those changes relate to other morphogenetic events such as cell division, tissue rearrangements or cell deformation. Here, we introduce the sea star embryo (*Patiria miniata*) as a model for spheroid epithelium dynamics, and we employ live imaging and machine learning segmentation algorithms to analyse cell and tissue shapes with respect to cell division and tissue compaction.

*P. miniata* embryos are transparent (Arnone et al., 2015; Meyer and Hinman, 2022; Newman, 1922), develop freely in sea water (Arnone et al., 2015; Meyer and Hinman, 2022; Newman, 1922) and can be imaged live for extended periods of time (Barone and Lyons, 2022; Perillo et al., 2023; Swartz et al., 2021). When fertilisation occurs, the fertilisation envelope is raised, the zygote undergoes holoblastic cleavage and then develops into an approximately spherical blastula (Arnone et al., 2015; Barone et al., 2022; Dan-Sohkawa, 1976; Kominami, 1983), with a blastocoel encircled by a monolayered epithelium (Arnone et al., 2015; Barone and Lyons, 2022; Dan-Sohkawa, 1976; Kominami, 1983). Interestingly, during cleavage stages sea star cells adhere loosely to each other and line up against the fertilisation envelope, occupying all available space (Barone et al., 2022; Barone and Lyons, 2022; Maruyama and Shinoda, 1990). At these stages, the epithelium is not sealed, there are openings between the cells through which fluid can flow (Barone and Lyons, 2022; Dan-Sohkawa and Satoh, 1978). However, cell division planes are perpendicular to the epithelial surface, so that the tissue becomes thinner and expands laterally after each round of synchronous cell division. Eventually, at around the 512-cell stage, closure is achieved and cells become progressively more compacted, i.e. more tightly packed together. (Barone and Lyons, 2022; Dan-Sohkawa and Satoh, 1978). Concomitantly, cell fate specification takes place, as different domains of the embryo are established: vegetal pole cells inherit maternal determinants that induce mesendodermal cell fates, while animal cells will develop into neuroectodermal cell types (Barone et al., 2022; Cheatle Jarvela et al., 2016; Maruyama and Shinoda, 1990; Nakajima et al., 2004; Swartz et al., 2021; Yankura et al., 2013; Zheng et al., 2022). Throughout cleavage and early blastula stages the embryonic shape remains approximately spherical (Barone and Lyons, 2022; Dan-Sohkawa, 1976).

Therefore, the sea star embryo is a dynamic spheroidal epithelium, undergoing sealing, while cells within it divide and differentiate. The combination of live imaging of sea star embryo development, detailed image analysis and computational modelling, allows us to ask whether, and how, morphogenesis and 3D epithelial packing are coupled.

## RESULTS

### Scutoids are induced upon global increase in cell density

To identify factors that couple cell proliferation and epithelial packing, we analysed the shape and 3D connectivity of cells in sea star embryos (*Patiria miniata*), which develop into approximately spherical monolayered blastulae (**Fig. 1A-B, Movie 1**) (Arnone et al., 2015; Barone and Lyons, 2022). In the sea star embryo, early blastomeres adhere loosely to one another initially, with fluid flowing between the inside and outside of the embryo until about the 512-cell stage, when the epithelium closes to encircle the blastocoel (Barone et al., 2022; Barone and Lyons, 2022; Dan-Sohkawa and Satoh, 1978) (**Fig. 1A, Fig. S1, Movie 1**). Importantly, during blastulation, many of the cellular processes characteristic of dynamic epithelia can be readily observed: cells undergo several rounds of cell division reducing their volume, changing shape and cell-cell contact topology along the whole height of the cells (3D packing) (Farhadifar et al., 2007; Lemke and Nelson, 2021; Nelson, 2016). To determine how cellular characteristics vary during development we performed time-lapse imaging of wild-type (WT) sea star embryos expressing a membrane marker (mYFP) (**Fig. 1A-B**) and a nuclear marker (H2B-CFP) combined with a deep-learning based 3D segmentation program (**Fig. 1C**, **Fig. S2**, see **Materials and Methods**). To ensure accurate segmentation we limited our analysis to the portion of the imaged embryos with the highest signal/noise ratio (**Fig. 1C**, see **Materials and Methods**).

**Figure 1.**
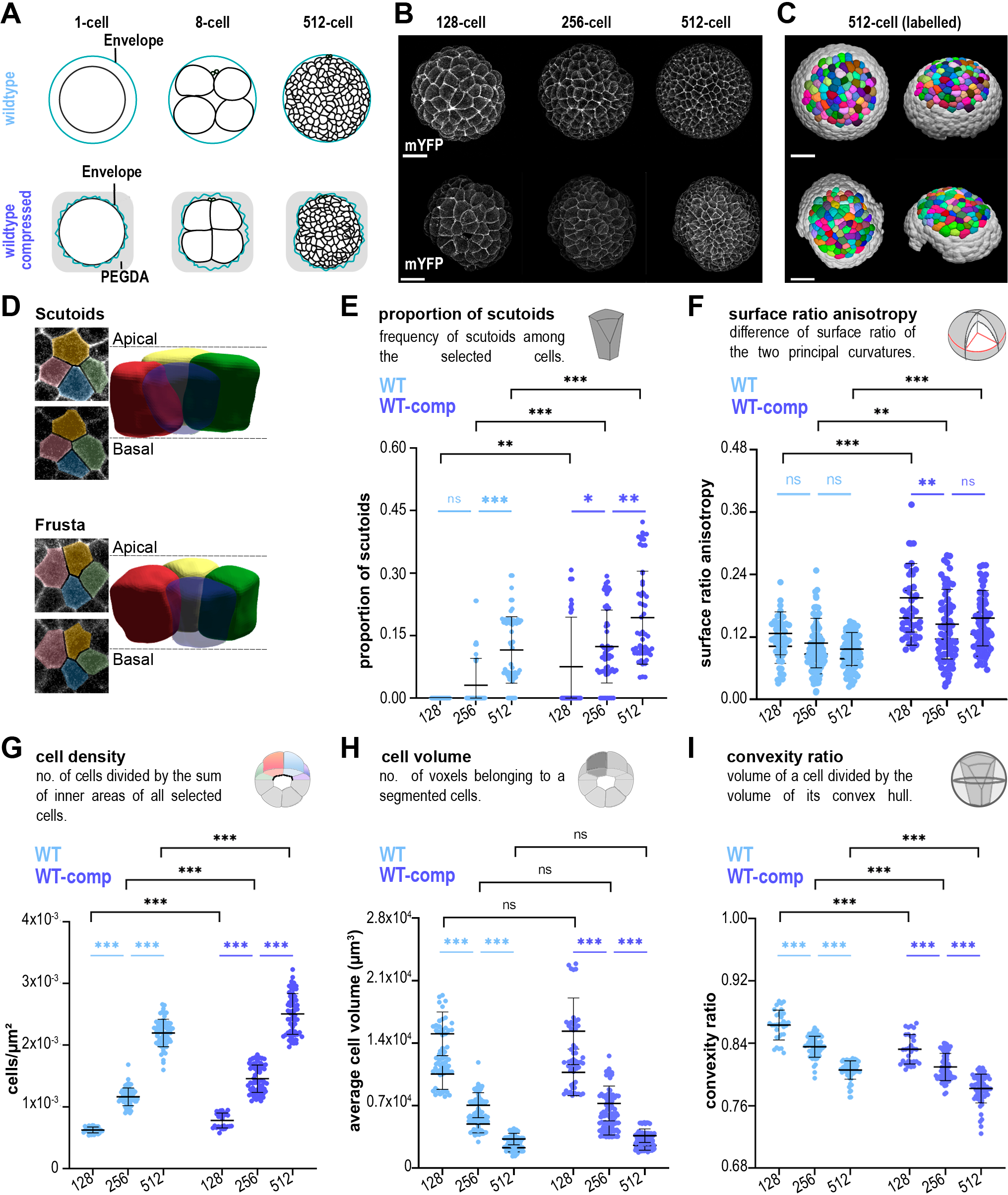
3D segmentation of sea star embryos over time: cell packing and topological analysis at cellular level. **A**) Schematic representation of WT (top) and WT-comp (bottom) sea star embryos. **B**) Maximum projections of a representative WT sea star embryo (top) or WT-comp embryo (bottom) expressing the membrane marker mYFP at 128-,256- and 512-cell stages. Scale bars, 50 μm. **C**) Computer rendering of the segmented sea star embryo at 512-cell stage from a frontal (left) and lateral (right) perspective. **D**) 3D representation of 4-cell motif with scutoid (top) or frusta conformations (bottom). The apical and basal z-slices of the motives are shown. Quantifications of average scutoid frequency (**E**), surface ratio anisotropy (**F**), cell density (**G**), cell volume (**H**) and cell convexity (**I**) are shown. WT: n=150 timepoints, 6 embryos, 4 experiments. WT-comp: n=150 timepoints, 6 embryos, 5 experiments for all panels except in F, where n=125 timepoints, 5 embryos, 4 experiments. Mean ± s.d. Statistical tests: Mann-Whitney tests with Bonferroni multiple comparisons correction (black) and Kruskal-Wallis tests with Dunn multiple comparisons correction, (blue) except in 1F where one-way ANOVA test with Tukey multiple comparison correction was used (light blue); ns: non-significant; *: p-value <0.05; **: p-value <0.01; ***: p-value <0.001.

The presence of “scutoidal” cells is a good indicator of changes in 3D packing, as scutoids are formed every time an AB-T1 occurs and the 3D connectivity increases (Gómez-Gálvez et al. 2022; Gómez-Gálvez et al. 2018; Okuda et al. 2019). Therefore, we set out to identify whether scutoids are formed in this dynamic spheroidal epithelium (**Fig. S3**), when they are formed, and which cell behaviours may be related to scutoid formation. We identified scutoids as cells with a different configuration of neighbours in the apical and basal sides (**Fig. 1D**) (Gómez-Gálvez et al., 2021a, 2018). Interestingly, we were not able to detect scutoids before the 256-cell stage by visual inspection. At the 256-cell stage, the epithelium started to seal, and then, we observed an increase in the frequency of scutoids at subsequent stages (**Fig. 1E**). Given that tissue geometry has been previously implicated in the appearance of scutoids (Gómez-Gálvez et al., 2018; Mughal et al., 2018), we asked whether curvature anisotropy changing over time explains the observed trend in scutoid formation. We calculated the surface ratio anisotropy of the region and found that it does not increase significantly over time (**Fig. 1F**). This result suggests that, in the context of an isotropic region of a curved tissue, factors other than curvature anisotropy are responsible for scutoid formation. To identify such factors, we analysed other cellular characteristics during embryonic development. First, we confirmed that cell density approximately doubled when the embryo advanced to a new stage (**Fig. 1G**), meanwhile cell volume was approximately halved after each round of cell division (**Fig. 1H**). When analysing cell shape changes, we found that cells became less convex (**Fig. 1I, Fig. S4**). Importantly, given that sea star cells are not protrusive (**Movie 1**), loss of convexity is likely attributable to how tightly cells are packed together. (**Fig S4**) and indicates increased cell compaction. These changes are expected given that the tissue expands laterally and becomes thinner, as shown by reduced cell height and tissue surface ratio (**Fig. S3**), which may result in increased compaction.

Taken together, these results suggest that AB-T1s may be facilitated by increased cell density and/or compaction. To further investigate these possibilities, we mechanically compressed and live-imaged sea star embryos by embedding them in a transparent viscous gel (Polyethylene Glycol Diacrylate hydrogel, PEGDA) at the 1-cell stage (**Fig. 1A-B, Fig. S1, Movie 2**). This procedure confines the embryo in a space that is smaller than usual, as normally embryonic cells occupy the entire space provided by the fertilisation envelope (**Fig. 1A**). We called this condition: WT compressed (WT-comp). Similarly to WT unperturbed embryos, we analysed the WT-comp at subsequent stages of development. We observed that surface ratio anisotropy did not increase (**Fig. 1F**), cell density doubled (**Fig. 1G**), cell volumes were halved (**Fig. 1H**), and cells became less convex (**Fig. 1I, Fig. S4**). However, the comparison between WT and WT-comp at each developmental stage highlighted interesting differences: compression induced a higher proportion of AB-T1s and their appearance at an earlier stage (128-cell in WT-comp vs 256-cell in WT) (**Fig. 1E**). Concomitantly, WT-comp embryos showed higher surface ratio anisotropy (**Fig. 1F**), higher cell density (**Fig. 1G**) and lower convexity (**Fig. 1I**) than WT embryos. Cell volume was the only feature not altered by the compression (**Fig. 1H**). Importantly, we have found that compression does not impact the timing of development and the synchronicity of cell divisions during the mitotic waves that occur between the 128- and 512-cell stages (**Fig S5**). These experimental results suggested that the dynamics of scutoid formation are linked to increase in cell density and tissue compaction.

### Regional differences in tissue compaction affect propensity of scutoid formation

During blastulation in the sea star embryo, several openings, located in different regions, close progressively; (**Fig. 2A-B, Fig. S6, Movies 1, 3**). We investigated this phenomenon in more detail by analysing separately the time-lapses where either the vegetal or the animal pole was imaged (**Fig. 2A, Fig. S6, Movies 1, 3,** see **Materials and Methods)**. We observed that closure is delayed on the animal pole: all openings are closed in vegetal poles of WT embryos by the 256-cell stage while they close only at the 512-cell stage in animal poles (**Fig. 2B, Fig. S6**). We then asked whether this asynchronous closure results in local differences in cell density and cell compaction that may induce scutoid formation. When we measured cell density, we found that it is not significantly different between animal and vegetal poles (**Fig. 2C**). However, using cell convexity as a measure of compaction, we found that vegetal cells had lower convexity than animal cells (**Fig. 2D**). Interestingly, scutoids appeared earlier and a higher proportion of cells acquired the scutoidal shape in the vegetal pole compared to the animal pole of WT embryos (**Fig. 2E**).

**Figure 2.**
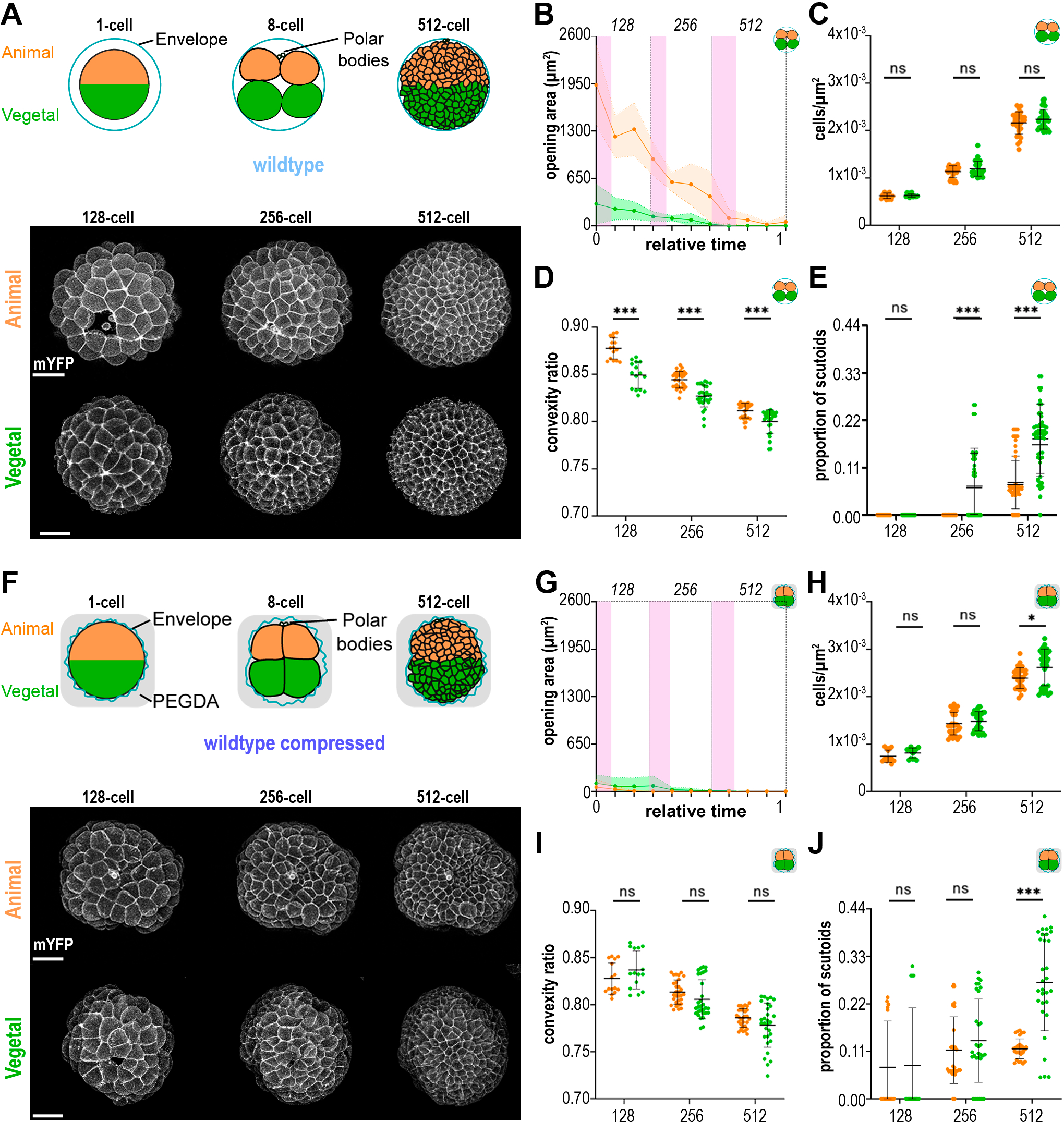
Asynchronous compaction and scutoid formation in the sea star embryo. **A**) (Top) Schematic representation of WT embryos highlighting animal and vegetal poles. (Bottom) Maximum projections of a representative animal pole and vegetal pole at 128-, 256- and 512-cell stages. Scale bars, 50 μm. Quantifications of opening areas (**B**), cell density (**C**), cell convexity (**D**) and proportion of scutoids (**E**) are shown. WT animal: n= 75, 3 embryos, 3 experiments. WT vegetal: n= 75, 3 embryos, 2 experiments. **F**) (Top) Schematic representation of WT-comp embryos highlighting animal and vegetal poles. (Bottom) Maximum projections of a representative animal pole and vegetal pole at 128-, 256- and 512-cell stages. Scale bars, 50 μm. Quantifications of opening areas (**G**), cell density (**H**), cell convexity (**I**) and proportion of scutoids (**J**) are shown. WT-comp animal: n= 75, 3 embryos, 2 experiments. WT-comp vegetal: n= 75, 3 embryos, 3 experiments. In (**B**) and (**I**) relative time 0 corresponds to the first cell division occurring at the 64-cell stage and relative time 1 corresponds to the first cell division occurring at the 512-cell stage. The pink areas denote the average duration of mitotic waves for each stage. Mean ± s.d. Statistical test: Mann-Whitney tests with Bonferroni multiple comparisons correction; ns: non-significant; ***: p-value <0.001.

These results suggested that, even though cell density was similar between poles, the presence of an opening allowed animal cells to remain slightly less compacted for longer, therefore delaying the formation of AB-T1s compared to the vegetal pole. In this scenario, the differences in the propensities for scutoid formation between animal and vegetal cells would be lost if the opening in the animal pole would be closed earlier. To test this hypothesis, we analysed compressed embryos (**Fig. 2F, Movies 2, 4**). We found that closure happens at the same time in animal and vegetal poles of WT-comp embryos, with openings being closed in both poles before the 256-cell stage (**Fig. 2G**). Cell density was similar between the two poles until the 512-cell stage, when it was higher in the vegetal pole compared to the animal pole (**Fig. 2H**). Moreover, there was no significant difference in convexity between animal and vegetal cells (**Fig. 2I**). Interestingly, there were no differences in the proportion of animal and vegetal cells acquiring the scutoid shape at the 128- and 256-cell stages, when both cell convexity and cell density were similar (**Fig. 2J**). At the 512-cell stage, however, when cell convexity was not significantly different but cell density was higher in the vegetal pole, a higher proportion of cells formed scutoids in the vegetal pole compared to the animal pole (**Fig. 2J**). While we observed differences in surface ratio anisotropy and cell volumes between animal and vegetal cells in a subset of the analysed stages, they did not correlate consistently with changes in the proportion of scutoids (**Fig. S4**).

Taken together, these results show that differences in compaction between regions of the embryo explain variation in the propensities for scutoid formation. Moreover, they show that compaction and cell density can independently affect 3D packing, as differences in the proportion of scutoids between regions with similar cell density can be explained by variation in compaction (WT) and, *viceversa*, differences between similarly compact regions can be explained by variation in cell density (WT-comp).

### Scutoid formation is temporally linked to cell divisions

Each stage of sea star development could be thought of as a steady state tissue characterised by a specific shape, compaction and cell density, which could alone account for differences in AB-T1 propensity. Alternatively, the increment of AB-T1s observed during development can also be due to dynamic factors. To discern between these two possibilities, we implemented spheroidal Voronoi models that mimic the shape and number of cells of each real embryo (as detailed in **Materials and Methods**). Briefly, we generated 3D Voronoi models for patches of epithelial cells corresponding to our experimental segmented cells. This means a construction that imitates the tissue with the same surface ratio anisotropy, cell density, and opening areas as in the embryos; but, importantly, without dynamic components such as cell proliferation (**Fig. 3A**). Then, we used the same principle to calculate the theoretical proportion of AB-T1s that should appear in both WT and WT-comp tissues. We observed a higher number of scutoids in virtual compressed tissues when comparing them with the WT constructions (**Fig. 3B**), showing that changes cell density and tissue geometry can influence the incidence of AB-T1s in the Voronoi model. However, we found that actual tissues presented a higher amount of scutoids than their corresponding Voronoi models (**Fig. 3B).** This result suggests that the appearance of scutoids is only partially due to changes in cell density/compaction altering the position of cells relative to each other and to the shape of the tissue, i.e. to geometry. Other factors, not accounted for in the model, can also induce AB-T1s.

**Figure 3.**
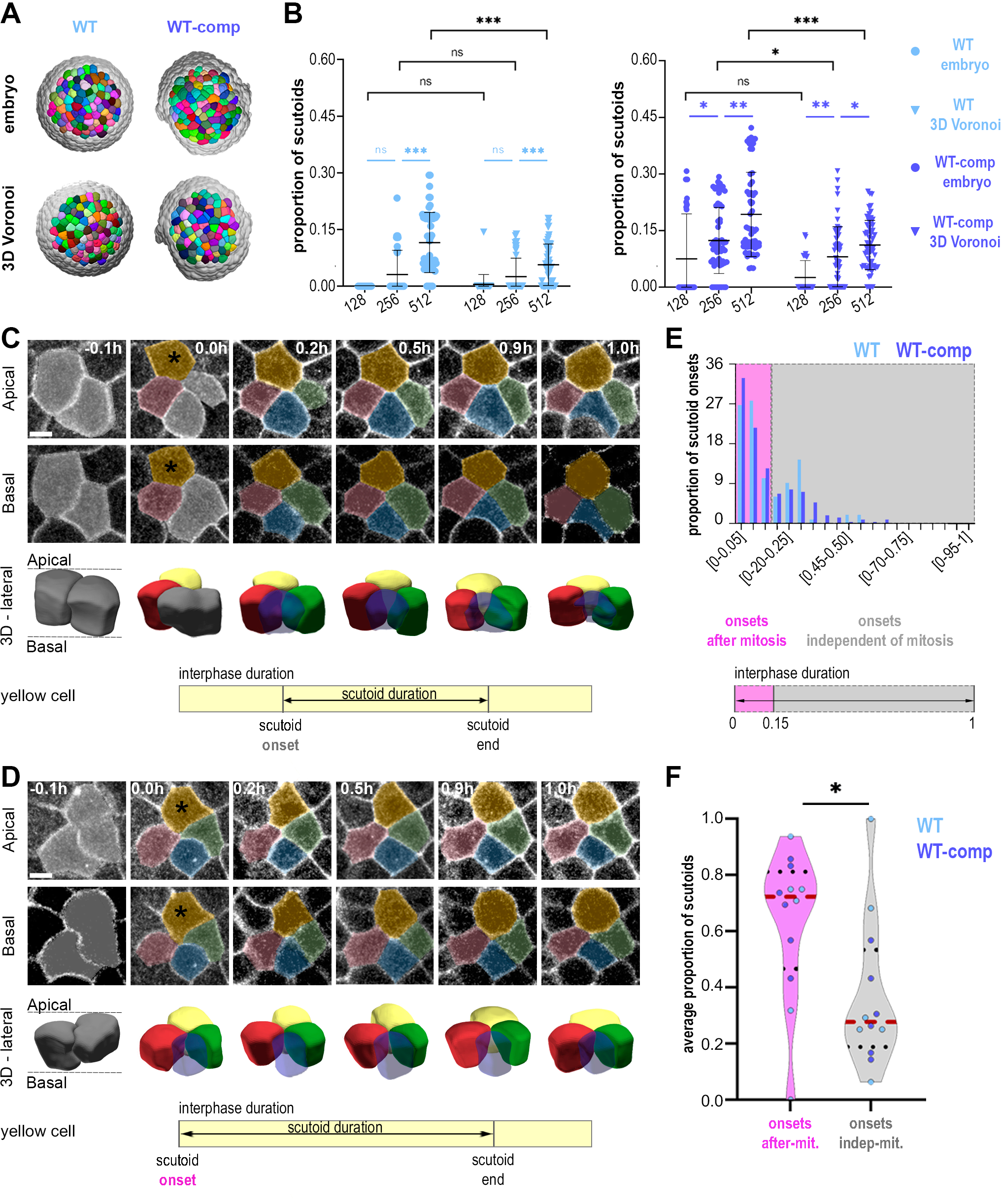
Single-cell tracking of scutoids relative to cell division. **A)** Computer rendering of 3D Voronoi models generated from WT and WT-comp embryos**. B)** Quantification of the proportion of scutoids. WT embryo: n=150 timepoints, 6 embryos, 4 experiments. WT 3D Voronoi: n=150 timepoints. WT-comp embryo: n=150 timepoints, 6 embryos, 5 experiments. WT-comp 3D Voronoi: n=150 timepoints. Mean ± s.d. Statistical tests: Mann-Whitney tests (black) and Kruskal-Wallis tests (blue) with Bonferroni multiple comparisons correction; ns: non-significant; *: p-value <0.05; **: p<0.01; ***: p-value <0.001. **C, D**) Representative scutoids whose onset is classified as independent (**C**) or after (**D**) mitosis Scale bars, 10 μm. Quantifications of the proportion of scutoids with onset after mitosis in the whole dataset (**E**) or per embryo (**F**) are shown. WT: 97 scutoids, 6 embryos, 4 experiments. WT-comp: 207 scutoids, 6 embryos, 5 experiments. Mean (red dotted line) ± s.d (black dotted lines). Statistical test: two-tailed Student t-test. *: p-value <0.05.

In the sea star embryo, rounds of synchronous cell divisions are responsible for the increase in cell density. Therefore, we used the time-lapses of the developing embryos to explore the relation between the formation of scutoids and cell division. We tracked individual cells over time and asked whether the formation of AB-T1s is temporally linked to cell division events within the epithelium (**Fig. 3C-F**). We analysed embryos between 128- and 512-cell stages and, for each scutoid detected, we tracked the cell during the whole interphase, and recorded when that cell acquired a scutoidal shape (scutoid onset) and when the AB-T1 transition was resolved (scutoid end) (**Fig. 3C-D**). In order to compare the timing of scutoid formation and duration between different developmental stages and different embryos, we normalised measurements over interphase duration, defined as the time between the end of cytokinesis and mitotic rounding marking the beginning of the following cell division (**Fig. S7**, **Materials and Methods**).

We tracked a total of 304 scutoid forming cells, of which 97 in WT embryos and 207 in WT-comp embryos (**Fig. 3E-F, Fig. S7**), as expected due to the higher frequency of scutoids observed upon compression (**Fig. 1E**). We found that the vast majority of scutoids in both conditions appeared shortly after cell division (**Fig. 3E**), with more than 60% of all scutoids being formed before 15% of the interphase time has elapsed, in both WT and WT-comp embryos (**Fig. 3E, Fig. S7**). This phenomenon was observed consistently across embryos, as in most cases (4 out of 6 WT embryos and 5 out of 6 WT-comp embryos) the proportion of scutoids with onset after mitosis - defined as before 15% of the interphase time - was higher than the proportion of scutoids with onset independent of mitosis - defined as after 15% of the interphase time - (**Fig. 3F**). Interestingly, in both WT and WT-comp embryos there was a steep decrease in scutoids onsets as the interphase progressed, with close to no scutoids being formed in the second half of the interphase (**Fig. 3E, Fig. S7**). Notably, we did not find a single cell that acquired the scutoidal shape twice within one interphase (**Fig. S7**). We observed no fluctuations in the position of the transition point along the apicobasal axis that resulted in the resolution and re-establishment of an AB-T1 among the same 4 cells. Instead, the apicobasal transition is formed rapidly and then the transition point moves slowly toward either the basal side or apical side until the AB-T1 is resolved and the scutoidal cells return to a frustum shape (**Fig 3C,D**).

Although scutoid onset was not altered, scutoid duration was increased in WT-comp embryos compared to WT embryos (**Fig. S7**). Not only average scutoid duration **(Fig. S7**) but also the proportion of scutoids that persisted through the second half of the interphase (**Fig. S7**) was higher in WT-comp embryos. The results from the experimental and computational experiments show a strong correlation between the formation of scutoids and cell division, suggesting a reorganisation of 3D cell packing as a response to maintain tissue homeostasis after the alteration induced by local increases of cell density.

## DISCUSSION

Elucidating the mechanisms involved in 3D packing is crucial to understand how tissues and organs form during animal embryogenesis (Gómez-Gálvez et al., 2021a; Lemke and Nelson, 2021). It has been previously proposed that AB-T1s are induced by i) curvature anisotropy (Gómez-Gálvez et al., 2018; Mughal et al., 2018) ii) steep curvature gradients that can lead to cell tilting (Lou et al., 2023; Rupprecht et al., 2017), and iii) cell migration and proliferation (Gómez-Gálvez et al., 2018). From a biophysics point of view, the formation of scutoids is clearly related to the surface tension parameters of the cells (Gómez-Gálvez et al., 2018; Mughal et al., 2018). In addition, Lou and colleagues propose that, in cylinder and ellipsoidal geometries, the interplay between mechanics (e.g. pressure, cell density and lateral tension) and cellular tilt is responsible for the appearance of neighbour rearrangements along the apico-basal direction (Lou et al., 2023). However, distinguishing the specific involvement of each factor affecting 3D packing has been challenging.

Here, we take advantage of the characteristic development of the sea star embryo, together with its amenability to live imaging and mechanical manipulations, combined with deep learning-based segmentation, to dissect the contributions of tissue compaction, cell density, and cell proliferation to the formation of AB-T1s and the consequent 3D tissue rearrangement. Importantly, our experiments and computational models discard a relevant contribution of curvature anisotropy, since the induction of scutoids does not consistently correlate with increased tissue surface ratio anisotropy (**Fig. 1E-F**). Instead, we find that scutoids form only once the epithelium is closed; then reorganisation of 3D packing is dependent on the level of tissue compaction and on local and global changes in cell density.

In the sea star embryo, cells organise in a monolayered epithelium that is initially leaky. It is only with subsequent rounds of oriented cell divisions that the epithelium becomes progressively more compact. Three aspects of epithelial morphogenesis contribute to compaction, i.e. increased cell density, epithelial closure, and cells becoming more tightly packed (**Fig. 4**).

**Figure 4.**
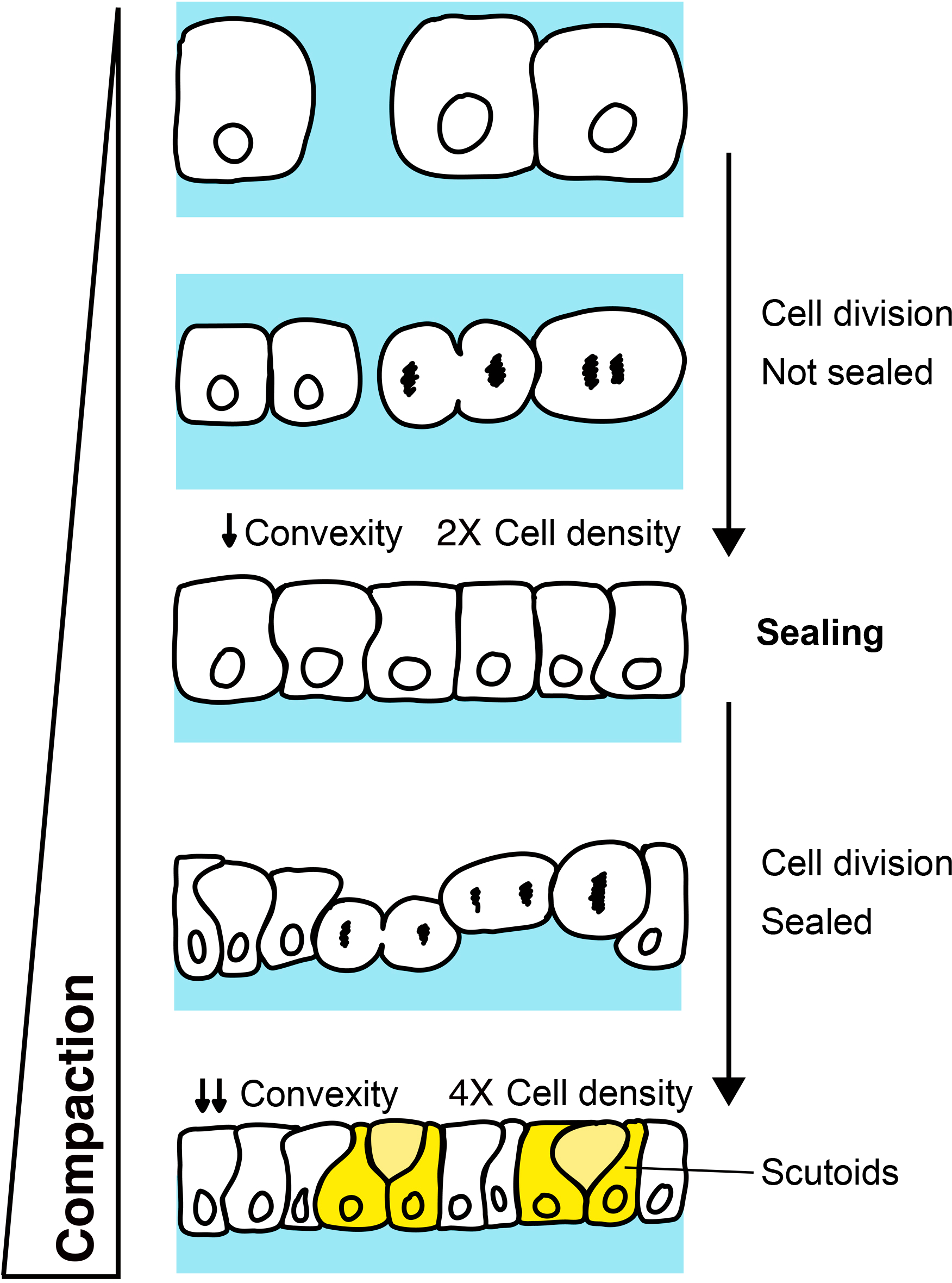
Relationship between compaction, cell density, convexity and the formation of scutoids. Schematic representation of the process of compaction in sea star embryos. Cell proliferation drives the lateral expansion of the epithelium via oriented cell divisions. Initially, cell divisions cause the reduction of interstitial space, until the epithelium is sealed. In this phase, when cells still have space to expand laterally, cell divisions do not drive the formation of AB-T1s. Once the epithelium is sealed, and cells are now confined, oriented cell divisions create lateral compression forces that result in lowered convexity and in cells adopting the scutoidal shape.

Cell proliferation occurs initially in a loosely packed epithelium and, only at later stages, in the context of a tightly packed tissue (**Fig. 4**). This phenomenon allows us to determine the relative contributions of proliferation and compaction to scutoid formation. In addition, the timing and extent of compaction is determined, at least in part, by how much space is available to the embryonic cells while the tissue expands laterally due to cell divisions. Therefore, we can perform two types of experiments: i) alter compaction by reducing such space, i.e. by mechanically compressing the embryo; and ii) use computational methods to model 3D packing in the absence of proliferation. Taking advantage of this model system and experimental approaches, we unravel the mechanisms underlying changes in cell packing at three different scales: whole-embryo (**Fig. 1**), embryonic domains (**Fig. 2**) and local proliferation (**Fig. 3**).

On a global scale, the analysis of the time-lapses shows that a significant amount of scutoids is formed only when cells are sufficiently compacted (starting at 256-cell stage), and then increase gradually in subsequent developmental stages (**Fig. 1E**). This implies that the increase in cell density *per se* is not sufficient to drive AB-T1s. Our results indicate that it is the coupling of increase in cell density with the space constraint of a compact epithelium that induces scutoid formation. This notion is supported by experimentally reducing the available space by compressing the embryo. In this case, sealing happens at the 128-cell stage, and so does the formation of scutoids.

On a second level, to address whether intrinsic factors may underlie scutoid formation, we took advantage of a peculiarity of the sea star embryo that we discovered in the course of our studies, i.e. sealing and compaction are heterogenous. In fact, epithelial closure is delayed in the animal pole with respect to the vegetal pole, resulting in lower compaction, even though cell density is similar (**Fig. 2B-C, Fig. S4**). The equally delayed appearance of scutoids in the animal pole indicates that tissue closure is necessary for the reorganisation of 3D epithelial packing (**Fig. 2A-B, E**). This conclusion is supported by our experimental setup: compression of the embryos synchronises closure, equalises compaction and results in synchronous induction of scutoids, which is anticipated to the 128-cell stage in both animal and vegetal poles of compressed embryos (**Fig. 2F-G, J**). In addition, our results show that when both poles are sealed and cells are equally compacted, higher cell density in the vegetal pole at the 512-cell stage results in higher proportion of scutoids (**Fig. 2H, J**). Therefore, this experiment shows that, once the epithelium is closed, cell density and compaction can affect 3D packing independently.

In this context, we propose that tissue compaction, measured here as reduced cell convexity, is a proxy for the pressure acting between cells within a tightly packed proliferating epithelium. Taken together, our results suggest that the extent of such pressure depends on the spatial constraints acting on the tissue (imposed either by the fertilisation envelope in WT embryos or by the embedding gel in compressed embryos) and, once the epithelium is closed, this pressure is combined with the pressure generated by increases in cell density. This, in turn, results in higher compaction and induces more AB-T1s. The balance of pressures acting on the tissue determines 3D packing.

Still, we find that the cells in the WT vegetal pole of the embryo are more compact and form more scutoids than cells in the animal pole at the 512-cell stage (**Fig. 2B, D-E**). Given that the external spatial constraints imposed by the fertilisation envelope are the same, these differences in compaction are probably due to differences in the material properties of the embryonic domains. Recent studies have shown that *in vivo* tissues undergo spatiotemporal transitions between fluid and solid states which present different properties, i.e. stiffness, cell motion and propensity of cellular rearrangements (T1 transitions) (Kuriyama et al., 2014; Mongera et al., 2018; Shellard and Mayor, 2023). Moreover, local fluidization or stiffening can change the balance of pressures acting on neighbouring tissues and ultimately drive morphogenesis (Barriga et al., 2018; Petridou et al., 2019). Conversely, cellular connectivity is also a determinant of tissue viscosity and stiffness (Petridou et al., 2021; Petridou and Heisenberg, 2019). Future studies, entailing direct measurements of cortical tension and viscosity in different embryonic domains (Sugimura, Lenne and Graner, 2016), will be needed to establish the relationship between AB-T1s and the physical properties of cells and tissues.

Finally, on a third level, the 3D epithelial packing comparison between real embryos and Voronoi models suggests that even accounting for tissue shape, tissue compaction and cell density is not sufficient to explain the high incidence of AB-T1s in a developing epithelium. Given that Voronoi models assume steady state conditions (Gómez-Gálvez et al., 2021b; Sánchez-Gutiérrez et al., 2016), while the sea star embryo is a highly dynamic proliferating tissue, we explore the role of local increase of cell density in scutoid formation. We find that the vast majority of cells acquiring a scutoidal shape do so shortly after cell division, both in WT and WT-comp embryos (**Fig. 3E-F**). We have shown that compressing embryos causes scutoids to appear earlier in development, and causes more cells to acquire the scutoidal shape. Yet, still, scutoids tend to form shortly after cell division in the experimental embryos (WT-comp) (**Fig. 3E-F**) at the same ratio as that in WT (**Fig. S7**). We think that this finding is in line with recent predictions from (Lou et al., 2023), who propose cell division as a source of pressure on cells’ lateral membranes that could induce scutoid formation. The tracking of individual cells after cell division shows that through the interphase the epithelium slowly accommodates and most scutoids are resolved into frusta (**Fig. S7**). In the compressed embryos, additional forces exerted on the cells exacerbate the phenomenon, causing the scutoidal shape to be maintained for longer periods of time (**Fig. S7**).

Altogether, we propose that, in the proliferating sea star embryo we have the combination of two phenomena: i) rounds of cell divisions where there is an increase in cell density and that lead to progressive tissue compaction and ii) the sudden local appearance of new cells after cell division, creating the need to cope with new neighbours (**Fig. 4**). Therefore, the induction of AB-T1s in the sea star embryo might be an efficient way to deal with the increasing pressures at local and global levels. Interestingly, it has been previously observed that the “scutoidal” shape is better than prisms at withstanding compression forces in an architectural context (Dhari and Patel, 2022). It is tempting to speculate that a side effect of scutoid formation in tissues undergoing morphogenesis is that they can better withstand compression.

Regarding future directions, our results add a new perspective on the regulation of 3D epithelial organisation and lay the foundation to identify the molecular mechanisms allowing cells to form and resolve AB-T1s in response to developmental changes.

## MATERIALS AND METHODS

### Animal husbandry

Adult *Patiria miniata* were purchased from Monterey Abalone Company (Monterey, CA) or South Coast Bio-Marine LLC (San Pedro, CA) and held in free flowing seawater aquaria at a temperature of 12-16°C. Sea star gametes were obtained as previously described (Hodin et al., 2019). Briefly, ovaries and spermogonia were dissected via a small incision on the ventral side of adults. Sperm was stored dry at 4°C while ovaries were fragmented to release oocytes in local filtered sea water (FSW). Maturation of released oocytes was induced by incubating for 1h at 16°C in 3 μM 1-Methyladenine (Fisher Scientific, 5142-22-3).

All embryos were raised in 0.22 μm - local filtered sea water (FSW) with the addition of 0.6 μg/ml Penicillin G sodium salt (Millipore Sigma, P3032) and 2 μg/ml Streptomycin sulfate salt (Millipore Sigma, S1277).

### mRNA injections

mRNAs were synthesised with the mMessage mMachine SP6 Transcription Kit (Invitrogen, AM1340). *Patiria miniata* immature oocytes were injected with mRNAs to label membranes (mYFP 100 ng/μl or mGFP, 400 ng/μl), and nuclei (H2B-CFP, 100 ng/μl or H2B-RFP, 400 ng/μl).

A subset of embryos were also injected with sp-ctnnb-RFP 800 ng/μl. Injected oocytes were incubated at 16°C overnight, activated and fertilised.

### Live imaging and embryo compression

*Patiria miniata* embryos expressing membrane and nuclear markers were mounted on a glass bottom dish (MatTek, P35G-1.5-14-C). No medium was used to immobilise the embryos: the glass bottom part of the dish was covered with a coverslip and sealed with vaseline. This creates a 500 μm deep chamber in which capillarity prevents the embryos from moving, until they develop cilia (Barone and Lyons, 2022). Additional FSW was added in the dish, to help with temperature control. Embryos were incubated until the 4-cell stage and then imaged on an inverted Leica Sp8 confocal microscope (20X objective, NA 0.7, 16°C controlled temperature).

Of the 6 embryos selected, 3 were oriented with the animal pole and 3 with the vegetal pole facing the objective. Orientation of the embryo was determined based on the position of the polar bodies and on the cleavage planes at 4- to 16-cell stages.

To achieve compression of developing embryos, 1-cell stage embryos were mounted in 3% PEGDA (EsiBio, GS700) in FSW. This restricts the embryos into a smaller space than normal, as they would otherwise occupy the entire fertilisation envelope. Embryos were otherwise imaged in the same way as control embryos.

The acquired timelapses were deconvolved using the Lightning module of the LeicaX software.

### Scutoid tracking

Scutoids were manually tracked using Fiji/ImageJ (Schindelin et al., 2012). Interphase for each cell forming a scutoid was defined as the time between the end of cytokinesis and cell rounding, which marks the beginning of the next cell division (**Fig. S7**). The time at which a cell adopted the scutoidal shape (scutoid onset) and for how long the cell maintained the scutoidal shape (scutoid duration) was recorded and normalised over interphase duration. Then, we classify scutoids into separate categories based on scutoid start times.

### Normalization of developmental time

To compare the duration of mitotic waves and dynamics of epithelial closure across embryos, developmental time was normalized by the time elapsed between the beginning of the 128-cell stage (relative time 0) and the end of 512-cell stage (relative time 1). Relative time 0 was defined as the time when the first cell of the 64-cell stage embryo had divided and relative time 1 as the time when the first cell of the 512-cell embryo had divided. Each stage was then split into a “mitotic wave” period, which ends when at least 50% of the cells have divided, and an interphase period, which ends when the first cell generated by the previous round of cell division divides again.

### 3D cell segmentation and tissue/cell feature analysis

For the automatic segmentation of 3D embryo stacks, we have followed a specific workflow pipeline adapted from a previously performed procedure called CartoCell (Andrés-San Román et al., 2023)

Training dataset was established from 3D Voronoi diagrams which was obtained combining the centroids of the cell nuclei of the sea star embryo as seeds and making masks from the cellular membranes to define the space to be filled. Custom Matlab scripts were used to calculate the nuclei centroids and the application VolumeSegmenter from Matlab were used to curate the membrane regions of the cells. The training dataset was composed of 14 time-points from 256-cells stage with 35 cells labelled of the same embryo development and was used as an input to a Deep Neural Network (DNN) presenting an architecture based on residual connections (3D ResU-Net) (Franco-Barranco et al., 2022).

Then, this model (M1) was tested with time points from other wildtype embryos even from other stages (128- and 512-). Finally the output of this model was used as input to another segmentation software, PlantSeg (Wolny et al., 2020), where through a watershed we obtain the labels of each of the cells that compose the stacks. Then, the mislabelled cells were checked using custom Matlab scripts. To improve predictions from wildtype and compressed embryos, a second model (M2) was trained by revising 150 stacks. In total, 300 stacks were processed from 12 embryo movies, with half belonging to wildtype embryos and the other half belonging to compressed ones. The segmented samples were obtained from three different cell cycles (128-, 256- and 512-cell stages). Specifically 30 time points were obtained from the earliest stage (5 per embryo) and 60 time points were obtained from each of the remaining two stages (10 per embryo and stage).

For each time point, we selected a subset of segmented cells for further analysis, i.e. the cells whose centroid lay within 30 μm from the embryo surface closest to the objective.

Note that custom made Matlab scripts were used to extract the following 4 characteristics:

- Proportion of scutoids: Frequency of scutoidal cells among the selected cells. We quantified the scutoidal cells marking the cells involved in AB-T1s, i.e., cells which exchange neighbours between apical and basal surfaces.
- Cell density of the region (cells/µm^2^): Ratio between the selected cells and the sum of inner basal areas they occupied.
- Average cell volume (µm^3^): mean volume of individual cells. The volume is the measurement of the number of voxels belonging to segmented cells.
- Cell convexity ratio: volume of the cell divided by the volume of the convex hull. The ratio ranges from 0 to 1, with values closer to 1 indicating a highly convex cell, and values closer to 0 indicating the opposite.
- Opening areas (µm^2^): Regions of the embryo not occupied by the cells. Once we extracted the full projection of the image stacks, we used FIJI’s polygon selection tool to quantify the sum of all opening areas in each sample. Note that for the following 4 characteristics, the samples segmented were 6 WT embryos and 5 WT-comp embryos.
- Embryo lengths (µm): To determine the dimensions of the embryo, the half embryo was estimated using the FIJI’s orthogonal view. The minor axis was measured by determining the distance from the uppermost region to the midpoint of the half embryo using FIJI’s straight line tool. The major axes of the embryo were measured on a 2D slice that approximately represented the midpoint of the embryo. These measurements were estimated using FIJI’s polygon selection tool.
- Tissue aspect ratio: Proportional relationship between the longest and shortest dimensions of a tissue. It is calculated by dividing the length of the longest dimension by the length of the shortest dimension. The aspect ratio provides information about the shape and elongation of an object. A value of 1 indicates a perfectly circular or square shape, while values greater than 1 suggest elongation along the longest dimension.
- Major axes lengths ratio: Proportional relationship between the two major axes of oblate spheroidal-shaped embryos.
- Average surface ratio anisotropy (or curvature anisotropy): Proportional relationship between the radii of curvature along the two main axes, *h* and *w* (transversal and longitudinal respectively) of the apical (*R_a_*) and basal (*R_b_*) surfaces. This relationship is characterised by the differences between the two surface ratios (Gómez-Gálvez et al., 2018). The formula to calculate the surface ratio anisotropy is as follows:

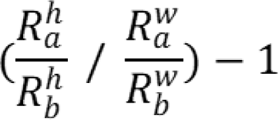

Where ℎ represents the axis of greatest curvature, *w* represents the axis of least curvature, *R_a_,* refers to the outer apical radius and *R_b_* is the inner basal radius. To obtain the value of these parameters, we measured the two principal curvatures for each centroid of the selected cells in the segmented region of both surfaces. By using the formulas to calculate the curvature of a coordinate in an ellipsoid (Bektas, n.d.), we obtained the values for the principal curvatures (*k_h_*,*k_w_*) and we computed the average value regarding the maximum and minimum radii of curvature 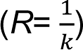 for both apical and basal surfaces. A surface ratio anisotropy of 0 indicates a tissue with an isotropic curvature, while values greater than 1 suggest a tissue with more anisotropic curvature.

### Voronoi constructions

We have used Matlab R2021a (Mathworks) as our computational tool to generate Voronoi constructions based on each imaged sea star embryo, maintaining similar geometrical characteristics of WT (n=150 time points, 6 embryos) and WT-comp datasets (n=150 time points, 6 embryos). These constructions were obtained by applying a 3D Voronoi algorithm to correctly mimic the epithelial packing (Voronoi 1908; Honda 1978). More specifically, we extracted the regions of the tissue that had been segmented from each imaged embryo in the form of binarized masks. These masks defined the bounding territory that the Voronoi cells could occupy in each construction. Then, we calculated the coordinates of the 3D centroids of all segmented cells which were then used as Voronoi seeds. The algorithm is based on tiling the space between these sets of Voronoi seeds by proximity, occupying the entirely binarized mask given in each construction (Voronoi 1908; Honda 1978).

Once the Voronoi construction was generated, similar to our approach done with the embryos, we selected a subset of Voronoi cells for further analysis, i.e. the cells whose centroid lay within 30 μm from the region that would correspond to the embryo surface closest to the objective.

### Delimitation of outer and inner layers and automatic detection of scutoids from segmented regions

The extraction of the outer and inner layers from the cellular segmented regions was performed using a custom MATLAB code (see *data availability*), traversing the segmented region along the Z-axis from top to bottom (outer) and bottom to top (inner) selecting the pixels corresponding to the first labelled cells encountered in both scans.

The method for extracting the proportion of scutoids from the Voronoi constructions was adapted from CartoCell (Andrés-San Román et al. 2023). After extracting the outer and inner layers, we quantified the number of neighbours for each cell on both surfaces. To identify neighbours, we dilated each cellular surface, identifying those labeled cells which overlapped with the dilation, thereby obtaining the sets of neighbours for a specific cell on each layer. Finally, we identify scutoids as cells that have different sets of neighbours on their apical and basal sides, as this situation can occur only in the case of an AB-T1.

### Statistical analysis

Statistical analyses were performed using GraphPad (Prism), as indicated in the figure captions. Shapiro-Wilk test was applied for normality determined our use of either standard Student t-test, ordinary one-way or two-way ANOVA tests (normally distributed data, equal variances) or non-parametric U-Mann-Whitney and Kruskal-Wallis tests (not normally distributed data). Bonferroni multiple comparisons correction was used to compare all features between WT and WT-comp conditions (**Fig. 1E-I**, **Fig. 3B** and **Fig. S3**) or between animal and vegetal locations (**Fig. 2B-E**, **G-J**, **Fig. S6**). Tukey multiple comparisons correction was used to compare WT surface ratio anisotropy over time (**Fig. 1F**, **Fig. S6**). Dunn multiple comparisons correction was used to compare all the other features over time (**Fig. 1E, G-I, Fig. 3B**, **Fig. S3, Fig. S6**). Student t-test was used to compare scutoids onsets after mitosis and independent of mitosis (**Fig. 3D**) and the average time elapsed between the beginning of the 128-cell stage and the end of 512-cell stage (**Fig. S5**). U-Mann-Whitney test was used to compare the average scutoids onset and duration (**Fig. S7**). The details of the statistical analyses for the different comparisons can be found on **Table S1**.

## Data availability

All data used in our analysis have been deposited at Mendeley Data and are publicly available at: https://doi.org/10.17632/45v8xcb5mp.

All original code used in our analysis is available at: https://github.com/ComplexOrganizationOfLivingMatter/seaStarProcessingSegmentation/releases/t ag/scutoidsProliferation2024.

Any additional information required to reanalyze the data reported in this paper is available from the lead contact upon request.

## Author contributions

Conceptualization: VB, AT, LME; Methodology: VB, AT, LME; Software: AT, JAS, JGG; Formal analysis: AT, VB; Investigation: VB, AT; Resources: DCL, LME, AH; Data curation: AT, VB; Writing— original draft: VB, AT; Writing—review & editing: VB, AT, DLC, LME; Visualization: AT, VB; Supervision: DCL, LME; Project administration: VB, LME; Funding acquisition: VB, DCL, LME.

## Acknowledgements

We would like to thank Pablo Vicente, Pedro Gómez-Gálvez and both the Lyons and Escudero labs for helpful discussions on the project and comments on previous versions of the manuscript. LME also wants to thank PIE-202120E047-Conexiones-Life network members for helpful input on the work.

## SUPPLEMENTARY FIGURES

**Figure S1.**
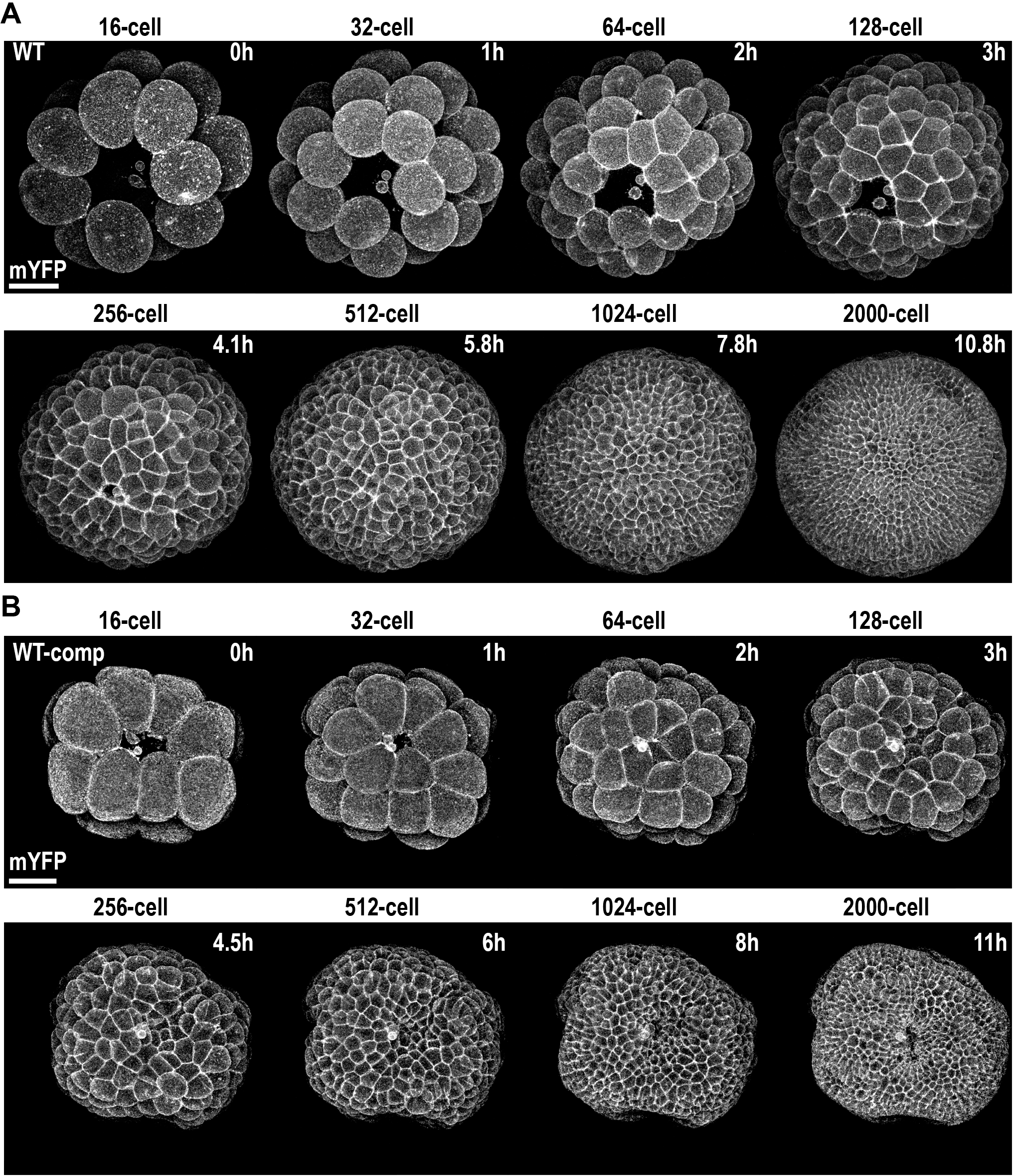
*Patiria miniata* embryo development. Maximum projections of a representative WT sea star embryo (**A**) or WT-comp embryo (**B**) expressing the membrane marker mYFP at different stages from 16-cells to 2000-cells. Scale bars, 50 μm.

**Figure S2.**
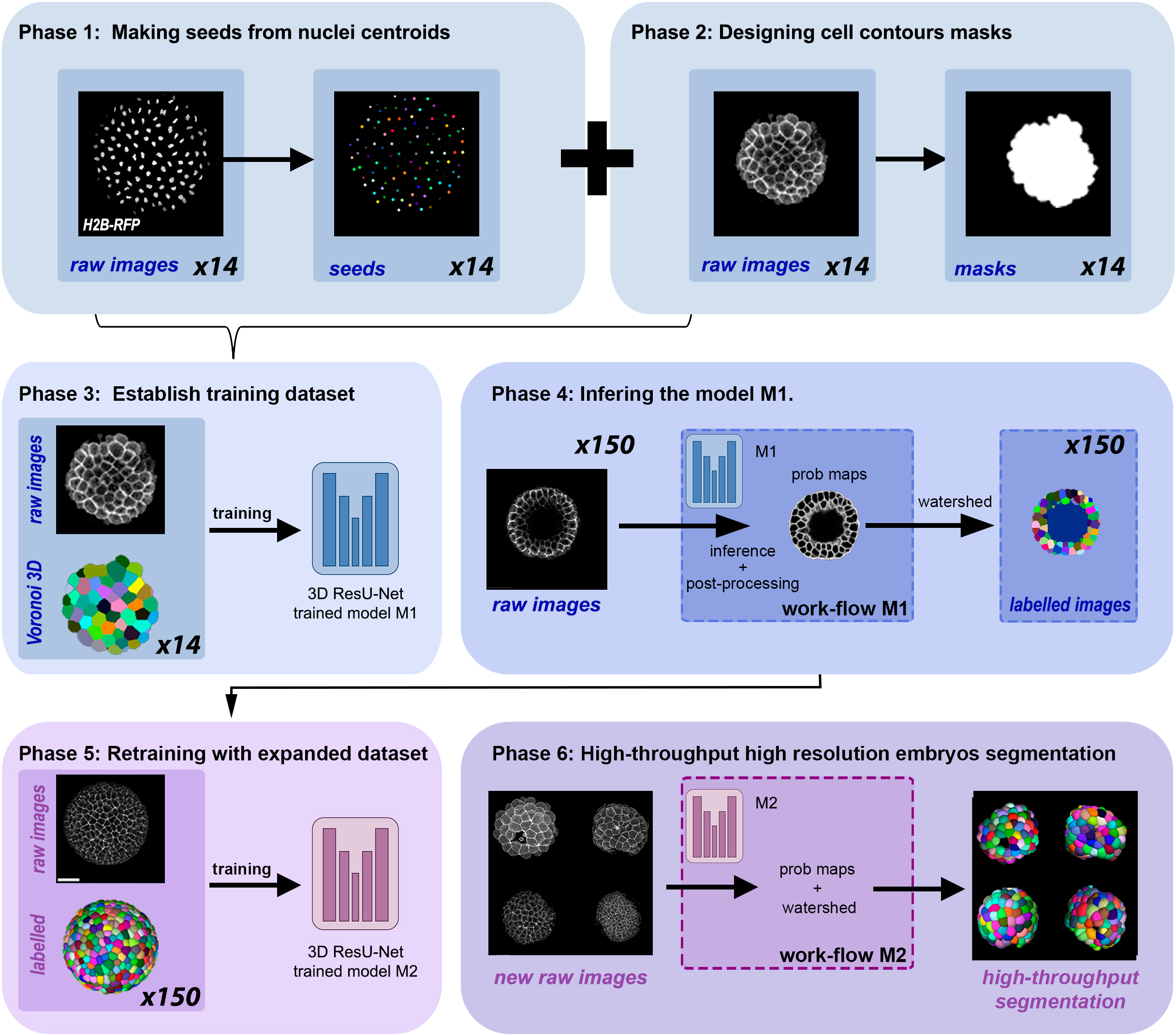
Deep learning-based 3D segmentation. Workflow scheme showing the different steps followed to segment sea star embryos.

**Figure S3.**
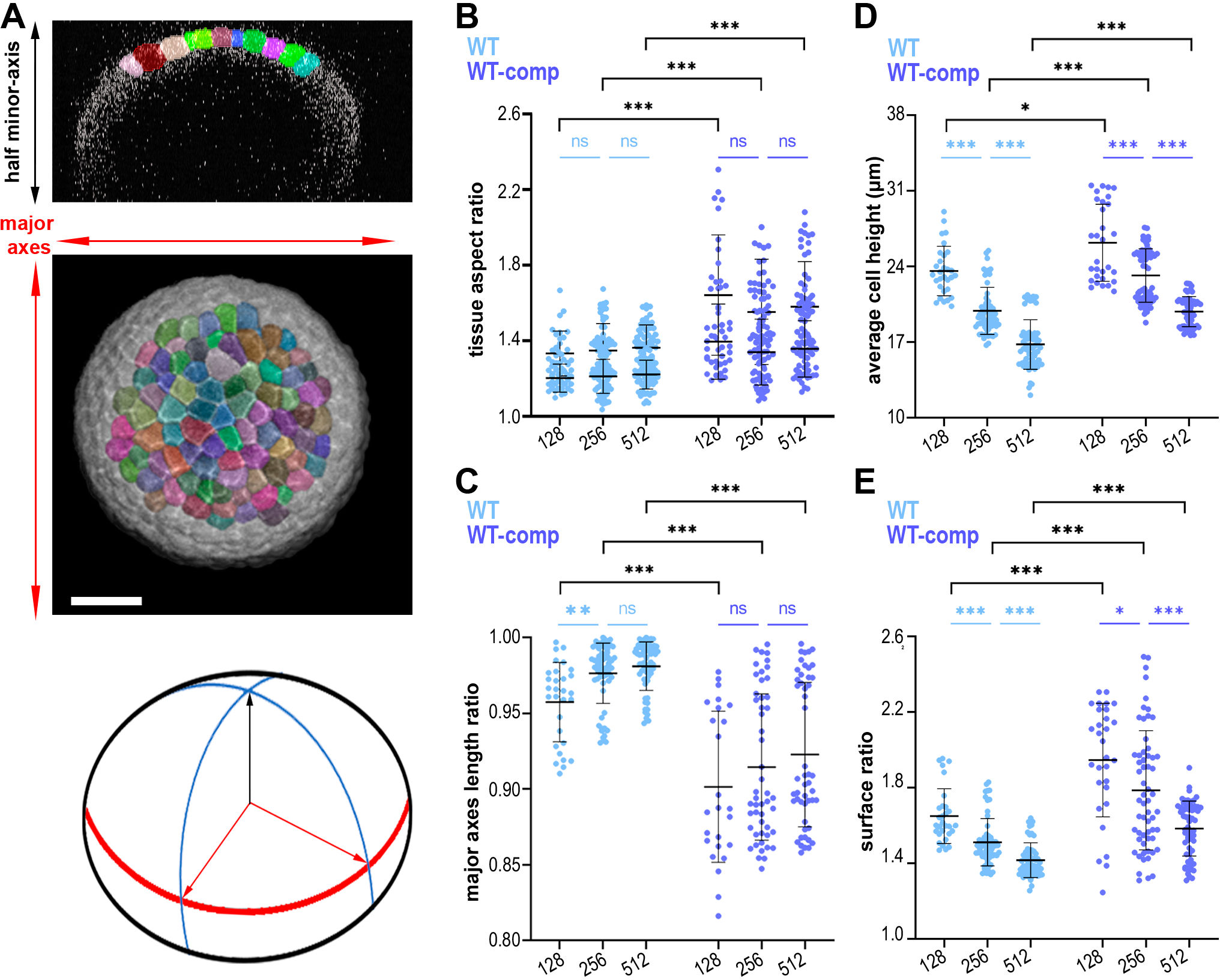
Shape of WT and WT-comp embryos at 128-, 256- and 512-cells. **A**) Orthogonal view (top) and maximum projection (centre) of a representative WT half embryo. (Bottom) Estimate of the complete shape. Scale bars, 50 μm. Quantification of the tissue aspect ratio (**B**) and major axes length ratio (**C**) of the whole embryos. Quantification of the tissue average cell height (**D**) and surface ratio (**E**) of the segmented region. WT: n=150 time points, 6 embryos, 4 experiments. WT-comp: n=125 time points, 5 embryos, 4 experiments for B-C panels, n=150 time points, 6 embryos, 5 experiments for D-E panels. Mean ± s.d. Statistical tests: Mann-Whitney tests with Bonferroni multiple comparisons correction (black) and Kruskal-Wallis tests (blue) with Dunn multiple comparisons correction; ns: non-significant; *: p-value <0.05; **: p-value <0.01 ***: p-value <0.001.

**Figure S4.**
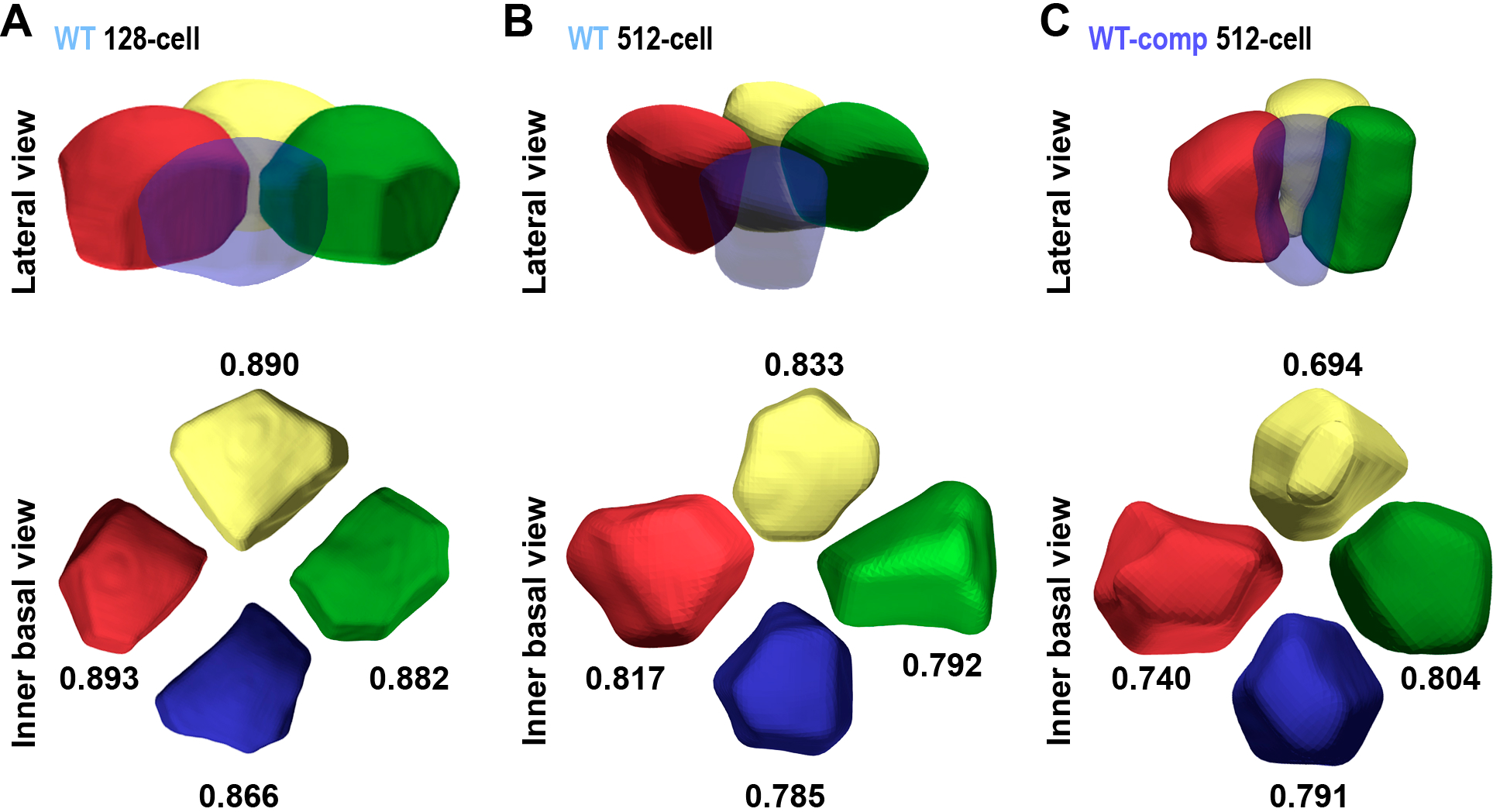
Cell compaction. 3D representation of 4-cell motives from a lateral (Top) and inner (Bottom) view from 128-cell WT embryo (**A**) 512-cell WT embryo (**B**) 128-cell stage (**C**) WT-comp embryo. In the inner view, the numbers displayed next to each cell indicate the individual value of the convexity ratio (**see Materials and methods**).

**Figure S5.**
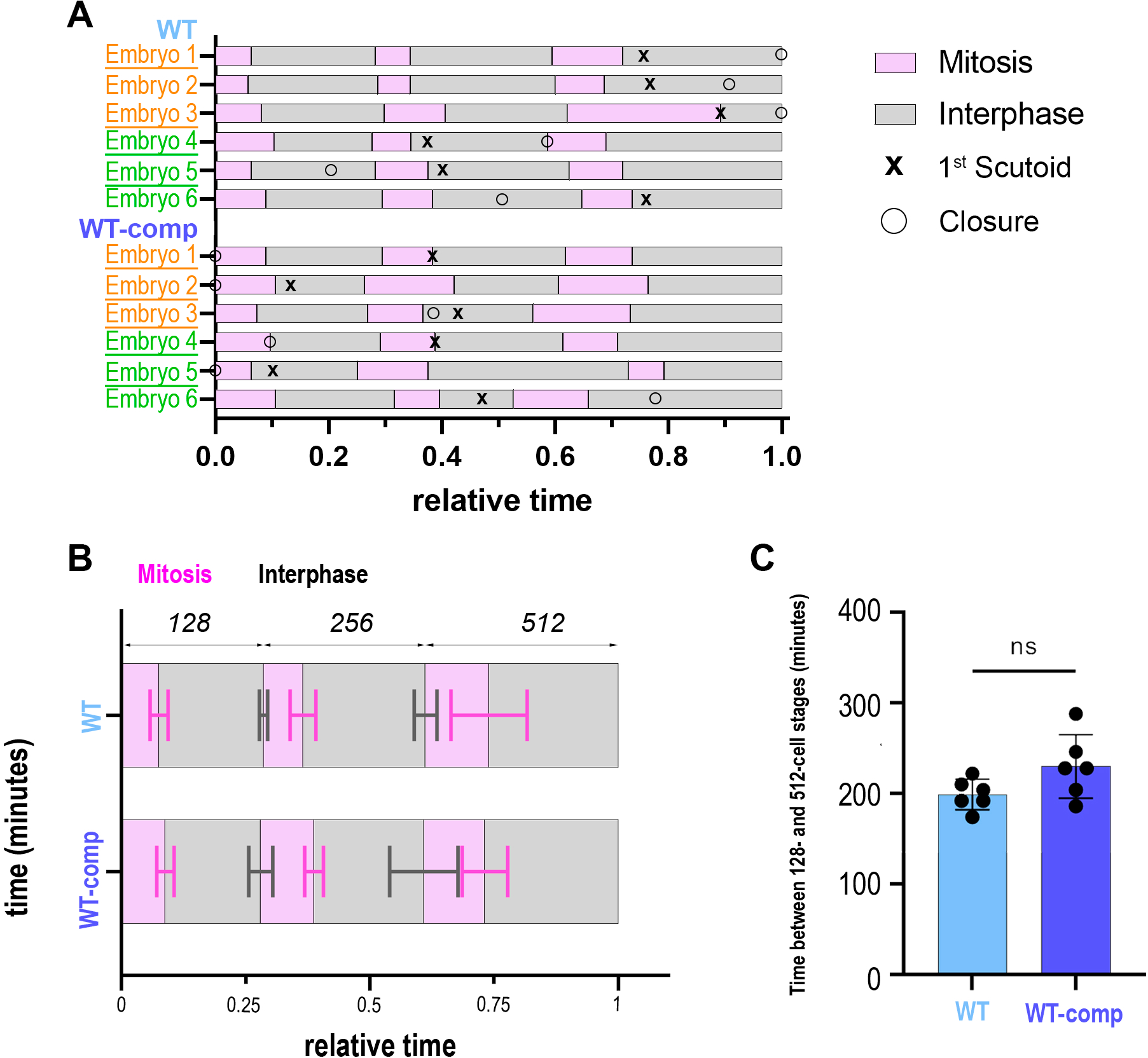
Single-cell tracking of division: analysis of cell proliferation rate per embryo. **A**) Quantification of mitosis and interphase time intervals per embryo showing the moment the tissue is completely sealed and the first cell adopting scutoidal shape. Quantification of the average time elapsed among each interphase and mitosis intervals (**B**) and between the beginning of the 128-cell stage and the end of 512-cell stage (**C**) in WT and WT-comp embryos. WT: n=150 time points, 6 embryos, 4 experiments. WT-comp: n=150 time points, 6 embryos, 5 experiments. Mean ± s.d. Statistical tests: two-tailed Student t-test; ns: non-significant.

**Figure S6.**
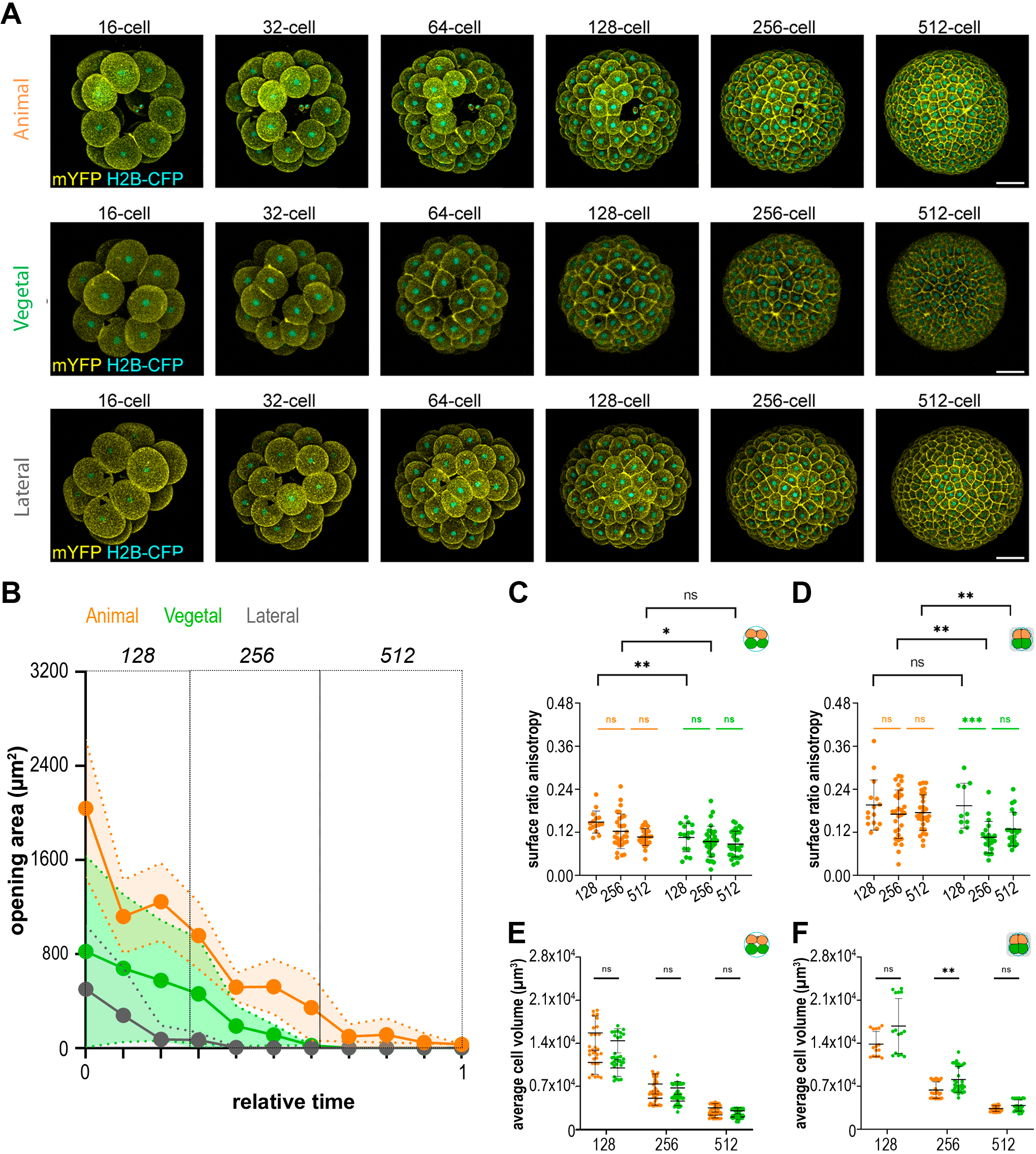
Opening areas and compaction. **A**) Maximum projections of representative WT sea star embryos expressing a membrane marker (mYFP) and nuclear marker (H2B-CFP) from an animal (top), vegetal (centre) or lateral (bottom) view. Scale bars, 50 μm. **B**) Quantification of opening areas in animal, vegetal and lateral regions. WT animal: n= 6 embryos, WT vegetal: n= 5 embryos; WT lateral: n= 4 embryos; 5 experiments. Quantification of surface ratio anisotropy in both regions of WT (**C**) and WT-comp (**D**). Quantification of cell volume in both regions of WT (**E**) and WT-comp (**F**). WT animal: n= 75 time points, 3 embryos, 3 experiments. WT vegetal: n= 75 time points, 3 embryos, 2 experiments. WT-comp animal: n= 75 time points, 3 embryos, 2 experiments. WT-comp vegetal: n= 50 time points, 2 embryos, 2 experiments for C-D panels, n= 75 time points, 3 embryos, 3 experiments for E-F panels. Mean ± s.d. Statistical tests Mann-Whitney tests with Bonferroni multiple comparisons correction (black) except in C where two-way ANOVA test with Bonferroni multiple comparisons correction was applied. Kruskal-Wallis tests with Dunn multiple comparisons correction (orange and green) except in C where one-way ANOVA tests with Tukey multiple comparisons correction was used; ns: non-significant; *: p-value <0.05; **: p-value <0.01; ***: p-value <0.001.

**Figure S7.**
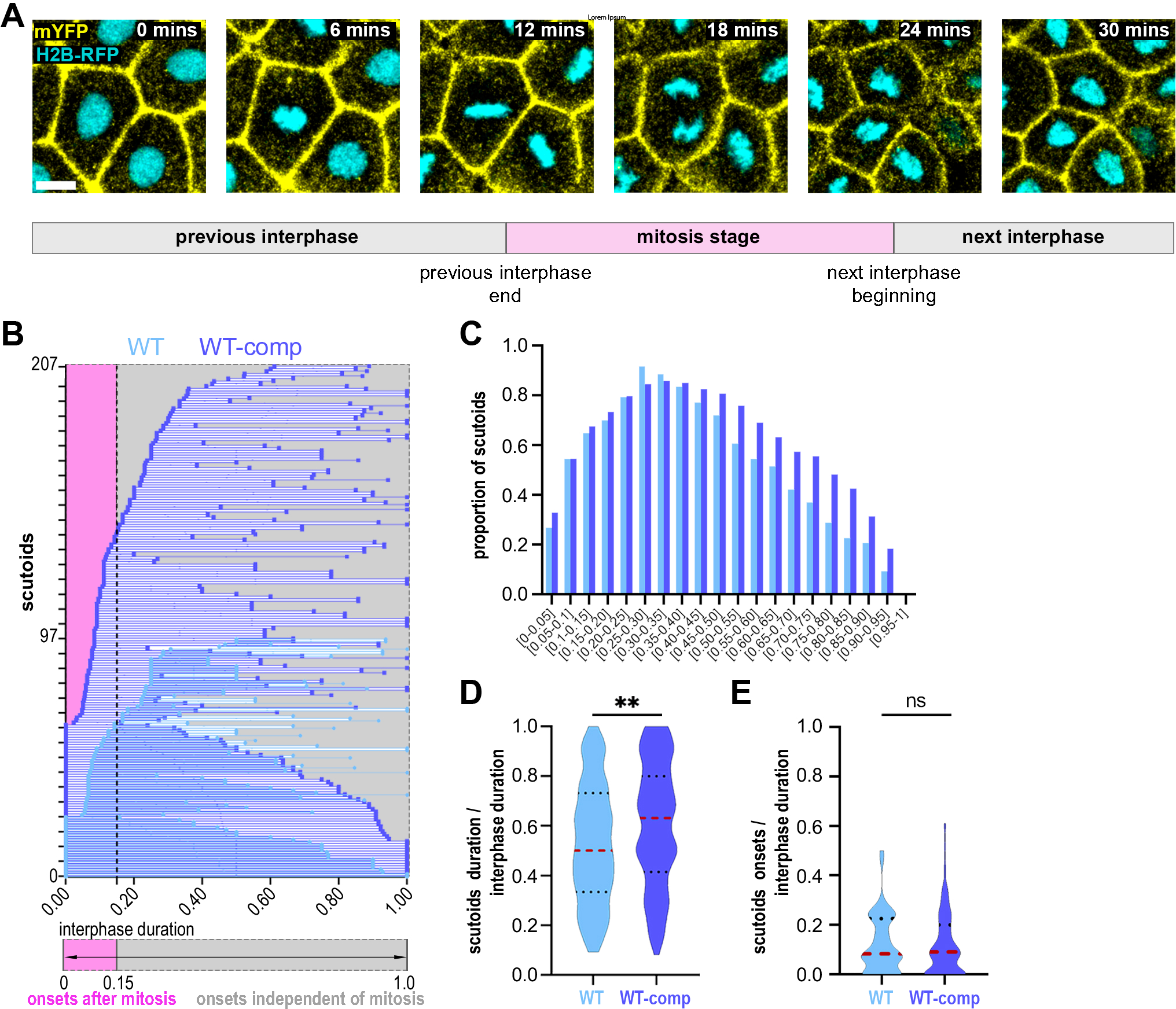
Tracking of scutoids over time. **A**) Slices of a representative sea star embryo showing how we establish the beginning and end of an interphase when tracking scutoids. The embryo is expressing the membrane marker mYFP and the nuclei marker H2B-RFP. Scale bars, 10 μm. Quantifications of the onsets and end of scutoids individually (**B**), the proportion of scutoids throughout the interphase (**C**), the average scutoids duration (**D**) and the average scutoids onset (**E**). WT: 97 scutoids, 6 embryos, 4 experiments and WT-comp: 207 scutoids, 6 embryos, 5 experiments. Mean (red dotted lines) ± s.d. (black dotted lines). Statistical test: Mann Whitney tests; ns: non-significant; **: p-value <0.01.

## SUPPLEMENTARY MOVIE LEGENDS

**Movie 1. *Patiria miniata* WT embryo, vegetal view.** Maximum projection of confocal time-lapse video of a WT embryo expressing a membrane marker (mYFP, yellow) and a nuclear marker (n-CFP, cyan) imaged between the 32- and 2000-cell stages. Vegetal view. Scale bars, 50 μm. Frame interval of 6 minutes, 7 fps.

**Movie 2. *Patiria miniata* WT-comp embryo, vegetal view.** Maximum projection of confocal time-lapse video of a WT-comp embryo expressing a membrane marker (mYFP, yellow) and a nuclear marker (nRFP, cyan) imaged between the 32- and 2000-cell stages. Vegetal view. Scale bars, 50 μm. Frame interval of 6 minutes, 7 fps.

**Movie 3. *Patiria miniata* WT embryo, animal view.** Maximum projection of confocal time-lapse video of a WT embryo expressing a membrane marker (mYFP, yellow) and a nuclear marker (n-RFP, cyan) imaged between the 32- and 2000-cell stages. Animal view (note the polar bodies). Scale bars, 50 μm. Frame interval of 6 minutes, 7 fps.

**Movie 4. *Patiria miniata* WT-comp embryo, animal view.** Maximum projection of confocal time-lapse video of a WT-comp embryo expressing a membrane marker (mYFP, yellow) and a nuclear marker (nRFP, cyan) imaged between the 32- and 2000-cell stages. Animal view (note the polar bodies). Scale bars, 50 μm. Frame interval of 6 minutes, 7 fps.

